# Stroke increases the expression of ACE2, the SARS-CoV-2 binding receptor, in murine lungs

**DOI:** 10.1101/2020.06.24.162941

**Authors:** Vikramjeet Singh, Alexander Beer, Andreas Kraus, Xiaoni Zhang, Jinhua Xue, Dirk M Hermann, Matthias Gunzer

## Abstract

**Background:** The newly emerged severe acute respiratory syndrome coronavirus (SARS-CoV-2) has caused a worldwide pandemic of human respiratory disease. Angiotensin-converting enzyme (ACE) 2 is the key receptor on lung epithelial cells to facilitate initial binding and infection of SARS-CoV-2. The binding to ACE2 is mediated via the spike glycoprotein present on the virus surface. Recent clinical data have demonstrated that patients suffering from stroke are particularly susceptible to severe courses of SARS-CoV-2 infection, thus forming a defined risk group. However, a mechanistic explanation for this finding is lacking. Sterile tissue injuries including stroke induce lymphocytopenia and systemic inflammation that might modulate the expression levels of surface proteins in distant organs. Whether systemic inflammation following stroke can specifically modulate ACE2 expression in the lung has not been investigated.

**Methods:** Mice were subjected to transient middle cerebral artery occlusion (MCAO) for 45 min and sacrificed after 24 h and 72 h for analysis of brain and lung tissues. Gene expression and protein levels of ACE2, ACE, IL-6 and IL1β were measured by quantitative PCR and Western blot, respectively. Immune cell populations in lymphoid organs were analyzed by flow cytometry.

**Results:** Strikingly, 24 h after stroke, we observed a substantial increase in the expression of ACE2 both on the transcriptional and protein levels in the lungs of MCAO mice compared to sham-operated mice. This increased expression persisted until day 3 after stroke. In addition, MCAO increased the expression of inflammatory cytokines IL-6 and IL-1β in the lungs. Higher gene expression of cytokines IL-6 and IL-1β was found in ischemic brain hemispheres and a reduced number of T-lymphocytes were present in the blood and spleen as an indicator of sterile tissue injury-induced immunosuppression.

**Conclusions:** We demonstrate significantly augmented ACE2 levels and inflammation in murine lungs after experimental stroke. These pre-clinical findings might explain the clinical observation that patients with pre-existing stroke represent a high-risk group for the development of severe SARS-CoV-2 infections. Our studies call for further investigations into the underlying signaling mechanisms and possible therapeutic interventions.

**Highlights:** Brain tissue injury increases ACE2 levels in the lungs

Brain injury induces pro-inflammatory cytokine expression in the lungs

Brain injury causes parenchymal inflammation and systemic lymphopenia

## 1. Introduction

Angiotensin-converting enzyme (ACE) 2 is present in mammalian tissues and plays an important role in the resolution of inflammation and cellular homeostasis under inflammatory conditions. The stimulation of ACE2 with specific activators is protective in specific diseases, such as brain injury induced by an ischemic stroke (Bennion et al., 2015; Mecca et al., 2011). However, recent data have demonstrated that SARS-CoV-2, the coronavirus causing COVID-19, utilizes ACE2 for entering into the epithelial cells (Hoffmann et al., 2020). The ensuing infection is accompanied by inflammatory lung injury and death and has caused a world-wide epidemic since its start at the end of 2019 (Pedersen and Ho, 2020).

Stroke-induced immune activation can affect multiple vital organs and augment the progression of specific inflammatory diseases. Previous findings have shown that the induction of stroke in murine models of brain ischemia can activate immune cells and promote the progression of inflammatory heart disease (Roth et al., 2018). Moreover, brain injury can modulate the functions of the intestine and its immune components, supporting the hypothesis of multi-organ failure after stroke (Singh et al., 2016a; Singh et al., 2018). In addition, stroke patients may present signs of severe immunosuppression and inflammation that often lead to hospital-acquired respiratory infections (Shi et al., 2018). A recent study by Austin et al. has demonstrated an increased number of mononuclear granulocytes in bronchoalveolar lavage fluid and higher IL-1β expression in lung tissue of mice that were subjected to ischemia-induced brain injury (Austin et al., 2019). In this respect, it is highly conspicuous that a plethora of clinical studies identified stroke as one of the most significant risk factors for a severe course of COVID-19 in humans (Bravi, 2020; Khan, 2020; Khawaja, 2020; Pigoga, 2020; Ssentongo, 2020). The reasons for this connection are currently enigmatic.

Hence, we hypothesized that brain injury-induced immunological alterations and systemic inflammatory conditions may enhance the expression of alveolar ACE2 and thereby, make stroke patients more susceptible to a lethal SARS-CoV-2 infection. At present, there is no data available demonstrating the dynamics of alveolar ACE2 and its functional role in lung inflammation after brain injury. Here, using an experimental murine model of stroke we found that focal cerebral ischemia indeed sharply increased the expression of ACE2 and pro-inflammatory cytokines IL-1β and IL-6 in the lungs.

## 2. Materials and methods

### 2.1 Animals

All animal experiments were performed following ethical guidelines and were approved by the local authorities (Landesamt für Natur, Umwelt und Verbraucherschutz Nordrhein-Westfalen, Essen, Germany). The C57BL/6Hsd wild type male mice are received from Envigo, Netherlands. In all experiments, mice of age between 8 and 10 weeks were used.

### 2.2 Mouse model of middle cerebral artery occlusion

Transient middle cerebral artery (MCA) occlusion (MCAO) was induced during anesthesia with 1% isoflurane in 100% oxygen. Mice were injected with the analgetic buprenorphine (0.1 mg/Kg, s.c.) and the anti-inflammatory drug carprofen (4-5 mg/Kg, s.c.). An eye ointment (bepanthen) was applied to avoid any harm to mouse eyes during the surgical procedures. A small incision was made between the ear and the eye to expose the temporal bone and a laser Doppler flow probe was attached to the skull above the core of the MCA territory. Mice were then placed in a supine position on a feedback-controlled heat pad and the midline neck region was exposed with a small incision. The common carotid artery (CCA) and left external carotid arteries were identified and ligated. A 2 mm silicon-coated filament (catalog #701912PKRe; Doccol) was inserted into the internal carotid artery to occlude the MCA. MCAO was verified by a stable reduction of blood flow to ≤20% of baseline that was observed on the laser Doppler flow device. Mouse wounds were sutured and mice inserted into a recovery box with a constant temperature of 37°C. After 45 min of MCAO, mice were anesthetized with isoflurane in 100% oxygen and the filament was removed for the reestablishment of the blood flow. Mouse wounds were again sutured and mice returned to their cages with free access to the food and water. Sham-operated mice underwent the same surgical protocol except the filament was inserted into the CCA and immediately removed. For experiments with 3 days survival, mice were daily injected with carprofen (4-5 mg/Kg, s.c.). The overall mortality rate of MCAO mice was ∼5% and that of sham mice was 0%. The exclusion criteria of experimental mice were as follows: inadequate MCAO (a reduction in blood flow to >20% of the baseline value), weight loss >20% of baseline body weight during the study, and spontaneous animal death.

### 2.3 Quantitative PCR

Mouse brain and lungs were processed to isolate total RNA using PureLink RNA Mini Kit (Invitrogen). An equal amount of total RNA was used to synthesize cDNA with a High-Capacity cDNA reverse transcription kit (ThermoFisher Scientific). Quantitative PCR was performed using a Q-Rex thermal cycler using the Rotor-Gene SYBR green PCR kit (Qiagen). Primer sequences were: ACE2 5” TGGGCAAACTCTATGCTG 3” and 5” TTCATTGGCTCCGTTTCTTA 3”; ACE 5” GTTCGTGGAAGAGTATGACCG 3” and 5” CCATTGAGCTTGGCGAGCTTG 3”; IL-6 5” CAG TTG CCT TCT TGG GAC TGA 3” and 5” GGG AGT GGT ATC CTC TGT GAA GTC 3”; IL-1β 5” TGTAATGAAAGACGGCACACC 3” and 5” TCTTCTTTGGGTATTGCTTGG 3”; GAPDH 5” AACTTTGGCATTGTGGAAGG 3” and 5” ACACATTGGGGGTAGGAACA 3”. The relative quantification was performed by the ΔΔCT method.

### 2.4 Western blotting

Mice were perfused with phosphate-buffered saline (PBS) and isolated lung tissues were prepared in lysis buffer (42 mM Tris–HCl, 1.3% sodium dodecyl sulfate (SDS), 6.5% glycerin and 2% protease and phosphatase inhibitors). The protein concentration of the samples was measured using the bicinchoninic acid protein assay kit (ThermoFisher Scientific). For immunoblotting, 20 μg of protein from each sample was subjected to SDS–polyacrylamide gel electrophoresis (PAGE) on a 10% gel under reducing conditions. Proteins were then transferred onto a polyvinylidene fluoride (PVDF) membrane (Millipore). After blocking with 5% milk solution, the membrane was incubated overnight at 4°C with the primary antibody. Primary antibodies used were goat anti-mouse ACE2 (AF3437, R and D Systems), goat anti-mouse ACE (AF1513, R&D Systems), and rabbit anti-mouse actin (4967, Cell Signaling). After extensive washing (three times for 15 min each in TBS containing 0.1% Tween 20), proteins were detected with horseradish peroxidase-coupled anti-goat or anti-rabbit antibody. Densitometry analysis was performed using ImageJ software (NIH, USA), and a β-actin control was used to confirm equal sample loading and normalization of the data.

### 2.5 Flow cytometry analysis

Mice were sacrificed and blood was collected via cardiac puncture and added to EDTA containing tubes. Mice were then perfused with PBS and spleens were collected in cold PBS. Single-cell suspensions were prepared by mincing the spleens in PBS and filtered through 70 μm cell filters. Blood and spleen samples were treated with RBC lysis buffer and washed in PBS. Total cell counts in samples were measured using an automated cell counter (Nexcelom Bioscience). An equal amount of cells from each sample was used to stain with antibodies purchased from Biolegend: CD45 FITC (30-F11), CD3 APC (17A2). Stained samples were measured on BD FACSAria™ flow cytometer and analyzed using FlowJo software (BD Biosciences).

### 2.6 Statistical analysis

Data were analyzed using GraphPad Prism version 6.0. All datasets were tested for normality using the Shapiro–Wilk normality test. The comparisons between two groups were analyzed using the Mann–Whitney *U* test. Differences with *p-*values≤0.05 were considered to be statistically significant.

## 3 Results

To understand the impact of stroke on lung ACE2 levels and inflammation, we utilized a well-established mouse model of focal cerebral ischemia, that is, transient MCAO. Induction of 45 min of MCAO in mice leads to focal brain infarcts that cover the striatum and overlying parietal cortex (Fig. 1a). Our results showed increased gene expression of pro-inflammatory cytokines IL-6 and IL-1β in the ipsilateral compared to contralateral brain hemispheres (Fig. 1b, c). In addition, mice subjected to stroke had a reduced number of circulating and splenic T lymphocytes which is an indicator of stroke-induced systemic immunosuppression (Fig. 1d, e).

**Figure 1.**
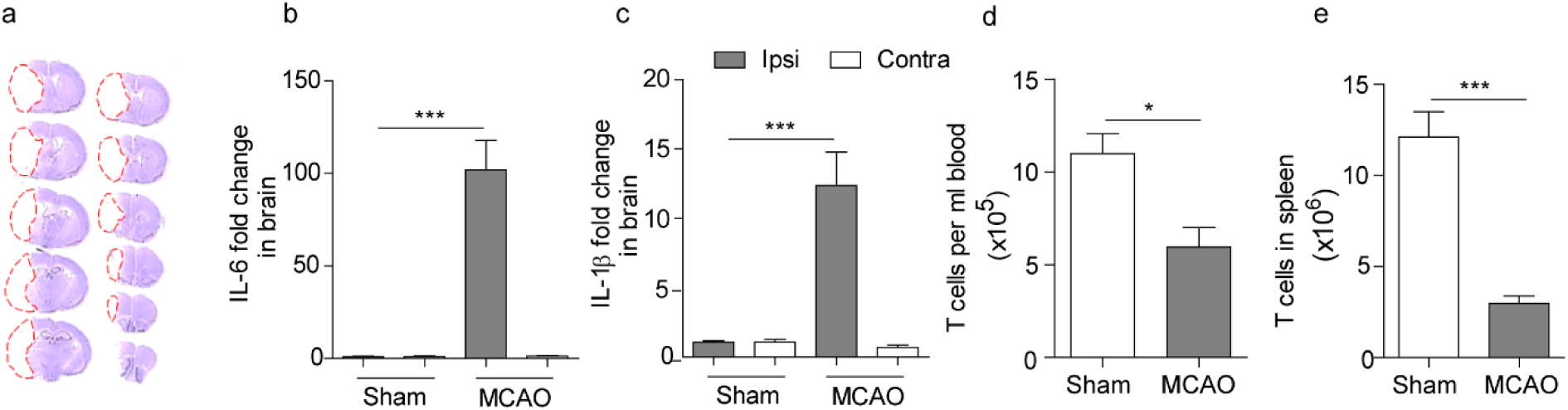
Sterile brain injury induces inflammation and lymphopenia. **a)** Brain serial sections stained with cresyl violet to visualize infarcts 24 h after transient MCAO. **b, c)** Increased gene expression of brain IL-6 and IL-1β in ischemic (Ipsi) hemispheres compared to non-ischemic hemispheres (Contra). **d, e)** Reduced number of T lymphocytes in blood and spleen of MCAO mice. Ipsi=ipsilateral; contra=contralateral. Data are means ± SEM. ***p<0.001, *p<0.05, Mann-Whitney U test, N=8 per group.

To test our hypothesis that stroke can increase the expression of ACE2 in the lungs, we performed quantitative PCR analysis on lung tissues 24 h after transient MCAO or sham surgery. Strikingly, the induction of transient MCAO significantly increased the level of ACE2 mRNA in the lungs compared to sham-operated mice (Fig. 2a). However, the expression of ACE mRNA was not changed between sham and transient MCAO (Fig. 2b). Interestingly, we also found an increased expression of the inflammatory cytokines IL-6 and IL-1β mRNAs in murine lungs after transient MCAO (Fig. 2c, d). These data suggest that stroke-induced peripheral signals can increase ACE2, but not ACE levels and generate an inflammatory milieu in the lungs of affected mice.

**Figure 2.**
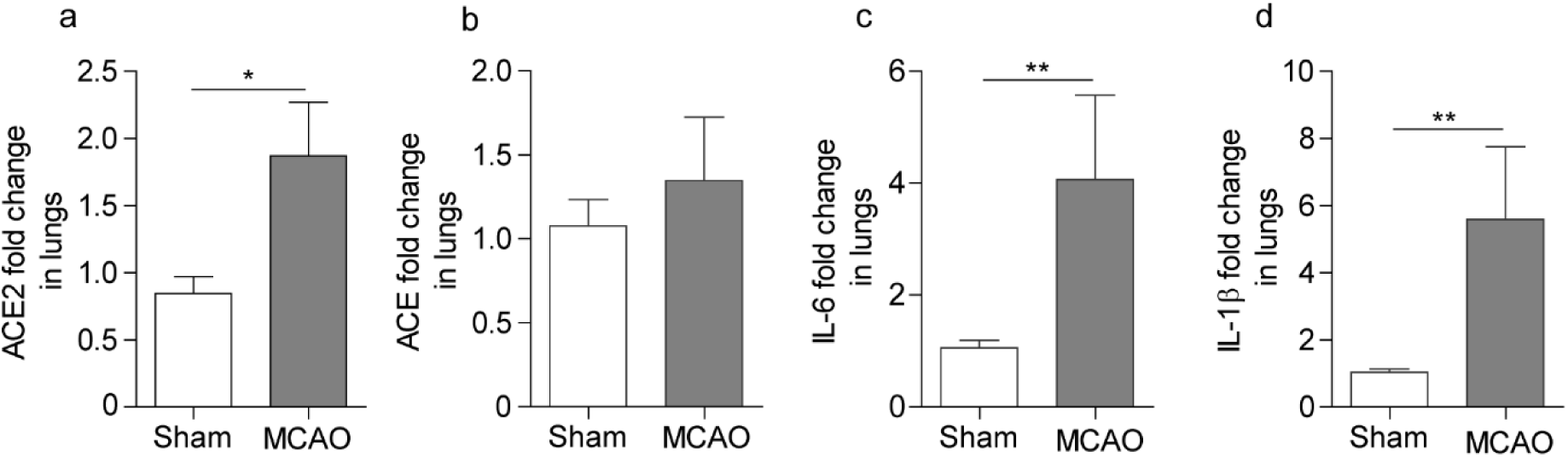
Brain tissue injury augments ACE2 mRNA expression and inflammation in the lungs. **a, b)** Increased gene expression of ACE2 but not ACE in the mice lungs 24 h after transient MCAO. **c, d)** Higher gene expression of IL-6 and IL-1β cytokines in the lungs of MCAO mice. Data are means ± SEM. **p<0.01, *p<0.05, Mann-Whitney U test, N=8 per group.

To verify that the increased mRNA expression was also mirrored on the level of expressed protein, we investigated the impact of transient MCAO on the abundance of ACE2 and ACE proteins in murine lungs by Western blotting. Indeed, we found that transient MCAO led to a 2-fold increase in ACE2 protein abundance compared to the sham controls 24 h post-ischemic-reperfusion injury. However, again, ACE protein abundance remained unchanged (Fig. 3a, b, c). We further extended our experiments to investigate the levels of ACE2 and ACE at a later time point of 72 h after transient MCAO or sham surgery. Again, we found increased levels of ACE2 but not ACE protein in lungs of MCAO mice compared to sham-operated mice (Fig. 3d, e). These results further indicate the long-lasting impact of systemic immune alterations and inflammation on ACE2 in lungs after sterile brain injury.

**Figure 3.**
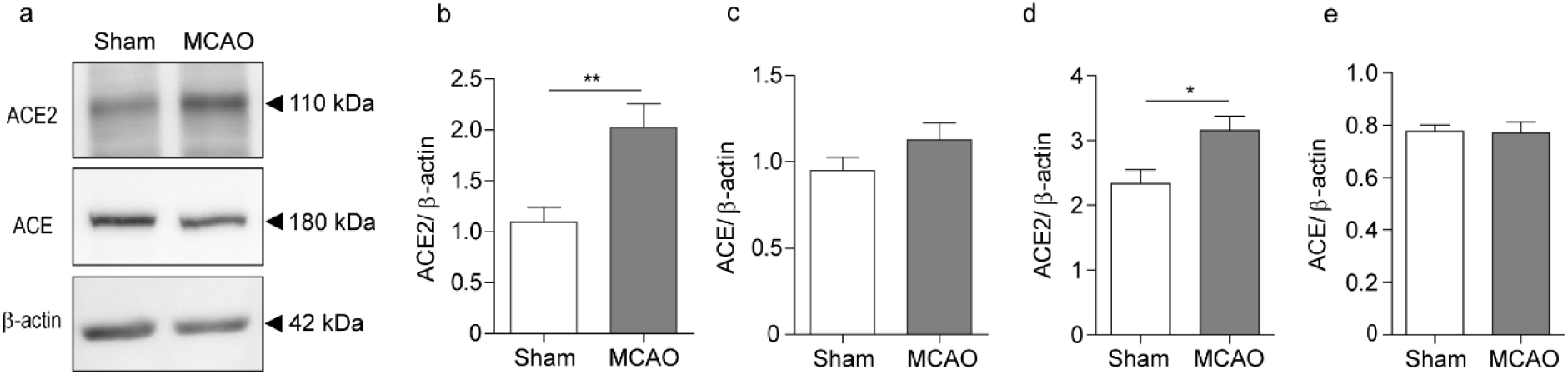
Brain injury increases ACE2 protein abundance in the lungs. **a)** Representative Western blots for lung ACE2, ACE, and β-actin protein in sham and transient MCAO mice. **b, c)** Increased levels of ACE2 but not ACE in the mice lungs 24 h and **d, e)** 72 h after MCAO. Data are means ± SEM. **p<0.01, *p<0.05, Mann-Whitney U test, N=6-8 per group.

## 4. Discussion

ACE and ACE2 are the key enzymes of the renin-angiotensin system which regulate blood pressure and salt-fluid balance in the body (Donoghue et al., 2000). ACE2 is a homolog of ACE and counterbalances pathways of inflammation in specific tissue injuries (Rodrigues Prestes et al., 2017). A higher expression of ACE2 is protective in lung injury, and the stimulation of this pathway with chemical activators has been shown to reduce LPS-induced lung edema and inflammatory tissue damage (Li et al., 2016). A recent study by Imai et al. suggested that the loss of ACE2 in a mouse model of acid-induced lung injury leads to the worsening of lung inflammation and increased lung edema (Imai et al., 2005).

ACE2 is utilized as a binding receptor by SARS-CoV2 for its entry into nasal and lung epithelial cells and the infection can induce acute respiratory distress syndrome (ARDS) in affected patients (Hoffmann et al., 2020; Ziegler et al., 2020). Based on the fact that SARS-CoV2 infection co-exists with reported cases of cardiac disease and stroke patients (Pan Zhai 2020; Shi et al., 2020), we hypothesized that brain injury may also increase ACE2 expression in the lungs. Indeed, our results suggest a rapid increase of ACE2 expression and pro-inflammatory cytokines in murine lungs after ischemic brain injury. The increase of ACE2 levels within 24 h of brain injury suggests fast kinetics of signaling to induce protein expression changes in the lungs. The molecular mechanisms underlying this response in the lungs after transient MCAO are currently unclear, but, based on our data and related studies of others increased post-stroke lung inflammation might be a possible explanation (Austin et al., 2019). Physiologically, the augmented ACE2 levels in inflamed lungs after brain ischemia may help to counterbalance the subsequent inflammatory lung injury (Imai et al., 2005). But in the presence of a virus that exploits this anti-inflammatory enzyme for cell entry and infection, this protective mechanism might prove fatal.

One of the clinical features of SARS-CoV-2-infected patients is the ensuing cytokine storm that leads to ARDS (Mehta et al., 2020). In the last months, several clinical studies have shown a higher serum level of cytokines such as IL-1β, IL-6 and TNF-α in virus-infected patients compared to the healthy controls (Conti et al., 2020; Han et al., 2020). Our results show that brain injury can also increase the expression of the pro-inflammatory cytokines IL-1β and IL-6 in the injured brain hemispheres and the lungs. In addition, sterile tissue injury can initiate the release of damage-associated molecular patterns (DAMPs) and thereby propagate parenchymal and systemic inflammation (Singh et al., 2016b). These DAMPs might serve as activation signals for lung epithelial and immune cells. However, which specific molecules modulate ACE2 expression in the lungs after stroke is an open question and requires further research.

The potential role of soluble ACE2 as a therapeutic target for SARS-CoV-2 infection is intensively discussed in the scientific community (Tang et al., 2020). Recent experimental data have demonstrated the effectiveness of recombinant human ACE2 in blocking SARS-CoV-2 binding to membrane ACE2 and thereby blocking virus invasion (Vanessa Monteil et al., 2020). The safety and use of soluble ACE2 therapy in humans has been successfully tested in a phase 1 clinical trial (Haschke et al., 2013) and is now examined in a clinical trial to treat COVID-19 patients (Clinicaltrials.gov #NCT04335136). In this respect, it is very important to note that decreased circulating levels of soluble ACE2 in stroke patients (Bennion et al., 2016) may further increase their susceptibility to SARS-CoV-2 infection in the light of increased levels of ACE2 in lung epithelial cells. Moreover, post-stroke immunosuppression is a key co-morbidity factor contributing to lung infections and leading to poor outcomes and increased mortality in affected individuals (Kalra et al., 2015). The mouse model of stroke used in our study also indicated a pronounced post-stroke lymphopenia similar to what is commonly observed after human stroke. Collectively, we suggest that increased ACE2 levels in lungs, systemic immunosuppression a reduced circulating ACE2 levels may act together to increase the prevalence of severe courses of COVID-19 in patients with preexisting brain injuries.

In conclusion, our results demonstrate that in the lungs of stroke mice ACE2 is rapidly upregulated and accompanied by increased inflammatory responses. The results described in this study may help to design new therapeutic strategies to reduce SARS-CoV-2 infections or ameliorate clinical courses of COVID-19 in stroke patients by targeting the ACE2 axis.

